# New functional identity of the essential inner membrane protein YejM: the cardiolipin translocator is also a metalloenzyme

**DOI:** 10.1101/2020.02.13.947838

**Authors:** Uma Gabale, Perla A. Peña Palomino, HyunAh Kim, Wenya Chen, Susanne Ressl

**Affiliations:** Indiana University Bloomington, Department of Molecular and Cellular Biochemistry, 212 S Hawthrone Dr, Bloomington, IN 47405, USA

## Abstract

Recent recurrent outbreaks of Gram-negative bacteria show the critical need to target essential bacterial mechanisms to fight the increase of antibiotic resistance. Pathogenic Gram-negative bacteria have developed several strategies to protect themselves against the host immune response and antibiotics. One strategy is to remodel the outer membrane where several genes are involved. *yejM* was discovered as an essential gene in *E. coli* and *S. typhimurium* that plays a critical role in their virulence by changing the outer membrane permeability by translocating and increasing the cardiolipin lipid concentration. How the inner membrane protein YejM with its periplasmic domain acts as a cardiolipin translocator remains unknown. Despite overwhelming structural similarity of the periplasmic domains of two YejM homologues with hydrolases like arylsulfatases, no enzymatic activity has been reported for YejM. Our studies reveal an intact active site with bound metal ions in the structure of YejM periplasmic domain. Furthermore, we show that YejM has a phosphatase activity that is dependent on the presence of magnesium ions and is linked to its cardiolipin translocation properties. Understanding the molecular mechanism by which YejM is involved in OM remodeling will help to identify a new drug target in the fight against the increased antibiotic resistance.

## Introduction

Indiscriminate use of antibiotics has resulted in a crisis of emergence of antibiotic resistant bacterial strains. The World Health Organization supports the larger scientific community in the statement that the world is running out of working antibiotics ^1^. Targeting specific mechanisms of antibiotic resistance is especially critical for Gram-negative bacteria such as *Escherichia coli* (*E. coli*) and *Salmonella typhimurium* (*S. typhimurium*) because of the recurrent outbreaks affecting society ^2–4^.

Pathogenic bacteria such as *E. coli* and *S. typhimurium* have developed several strategies to protect themselves against the host immune response and antibiotics. These mechanisms create antibiotic resistance allowing Gram-negative bacteria survival and replication ^5–7^. The bacterial cell membrane plays a critical role in limiting the effectiveness of antibiotics by acting as a barrier and preventing the diffusion of antibiotics and other harmful chemicals into the cell ^8,9^. Understanding the functional role of proteins and enzymes acting at the interface of the bacterial cell membrane is fundamental to address antibiotic resistance and address the effects of bacterial outbreaks.

The cell envelope of Gram-negative bacteria is made of an inner membrane (IM) and an outer membrane (OM), and the periplasmic space in between that contains a thin layer of peptidoglycan [2]. The OM is an asymmetric lipid bilayer; the inner leaflet is composed of phospholipids and the outer leaflet is made of lipopolysaccharides (LPS). Lipid A (endotoxin) is a component of LPS and a key virulence factor in endotoxin shock

^10^. The O-antigen polysaccharide that is linked to LPS is highly antigenic and has a striking flexibility and structural diversity ^11^. One strategy employed by Gram-negative bacteria to survive or hide from the host immune system is the modification of the lipid A moiety of LPS. This is facilitated by various inner membrane enzymes such as phosphoethanolamine (PEA) transferases, namely EptA and mobilized colistin resistance (MCR) family of proteins belonging to the alkaline phosphatase superfamily ^12–14^. They mediate the decoration of lipid A with PEA resulting in the OM resistance against cationic antimicrobial peptides (CAMPs) and lower affinity to Toll-like receptor 4 ^7,12^. Another strategy of OM remodeling is to increase the incorporation of cardiolipin, which is a crucial component of the OM that binds bacterial proteins during sporulation and cell division ^15,16^. Cardiolipin is synthesized at the inner membrane (IM), and cardiolipin trafficking in *Salmonella* to the OM was shown to be controlled by the two-component PhoPQ system ^17^. Increased levels of cardiolipin in the OM are found when bacteria experience stress, such as higher temperature (>=42°C)^18^, increased osmotic pressure ^19^, in stationary phase ^20^, as well as encountering host immune response during pathogenesis ^5^.

The essential gene *yejM* was discovered in *E. coli* and *S. typhimurium* ^21,22^. The homologue in *S. typhimurium* was termed *pbgA* (PhoPQ-barrier gene A) and plays a crucial role in increased cardiolipin levels in the OM ^22^, and linked to lipopolysaccharide (LPS) assembly defects in deletion mutants, specifically manifested by an increase in levels of lipid A core molecules, in a PhoPQ-dependent manner ^23^. *yejM* codes for an inner membrane protein with five predicted helical transmembrane domain (5TM), followed by a positively charged linker region and a C-terminal periplasmic domain (PD) (Figure 1a). YejM-PD is functionally predicted to be a sulfatase and it is fundamental to OM modifications and immune response. Mutants lacking the PD domain are viable, whereas deletion of the 5TM domain is lethal for Gram-negative bacteria ^21,22^. Importantly, *E. coli* strain LH530 lacking the YejM-PD showed an increased sensitivity to temperature and antibiotics such as vancomycin ^21^. However, *E. coli* strains lacking YejM were rescued by inactive or active forms of phosphopantetheinyl transferase (AcpT), that plays a minor role in lipid metabolism of *E. coli* K-12 ^24,25^. The YejM PD is also essential for the virulence of *S. typhimurium*; strains lacking YejM PD show no increase of cardiolipin in the OM, have increased permeability under highly PhoPQ activated environment, and fail to survive inside the host cells ^22^.

**Figure 1.**
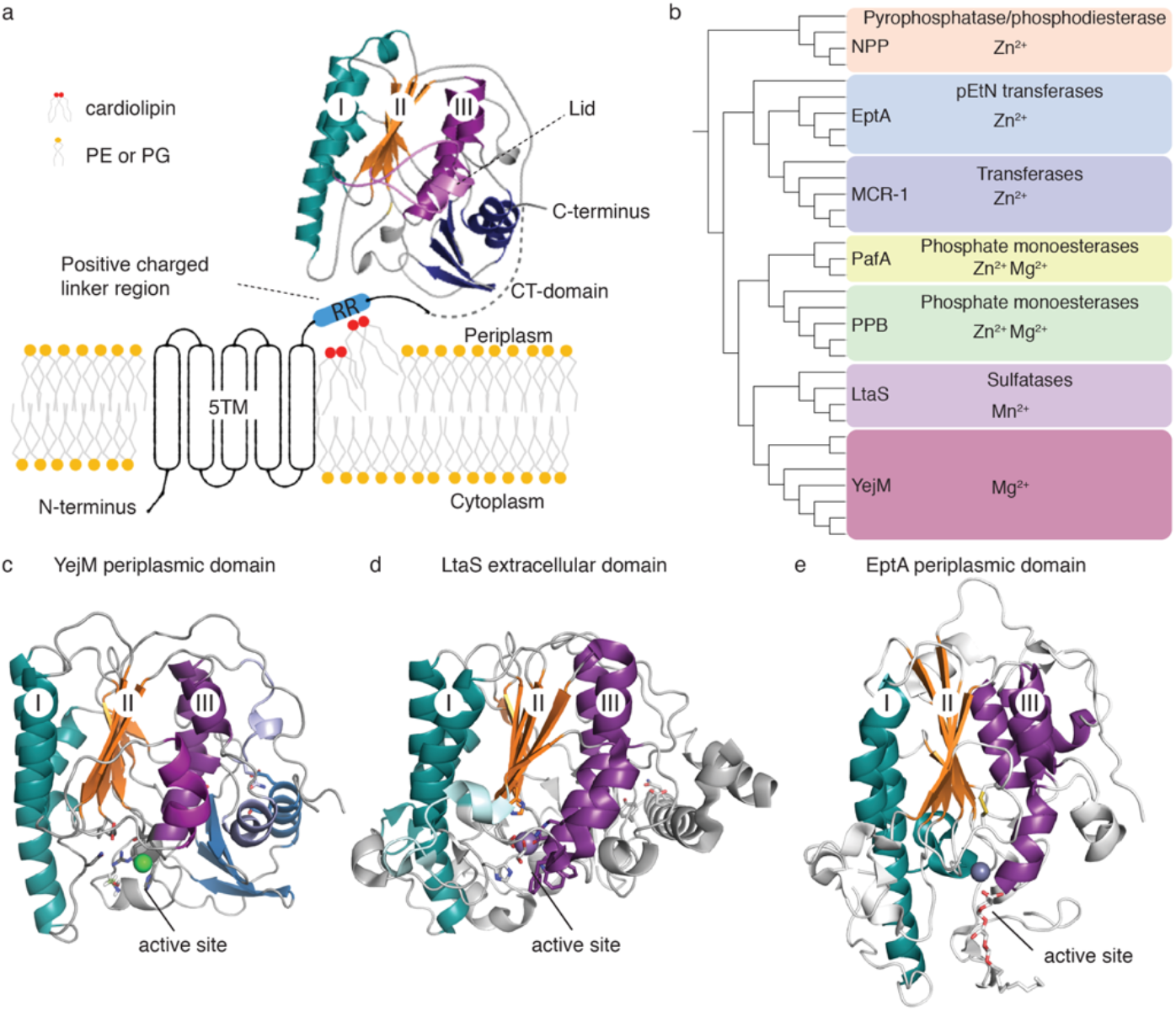
Overview of YejM structure, constructs, and phylogenetic connection of YejM to selected members of the larger hydrolase superfamily. **a** Overview of YejM structure. The N-terminal transmembrane domain containing 5 helices (5TM) and linker region with positively charged arginine residues (RR) are illustrated, cardiolipin lipids are indicated to bind to the RR linker region, PE and PG lipids are non-distinguished creating the membrane plane. The periplasmic domain (PD) is presented as our crystal structure of wildtype YejM PD with three layers: I turquoise, II orange, and III deep purple, indicated within the α/β hydrolase core domain. The lid in light purple *covers* the proposed hydrophobic cardiolipin binding pocket between layers II and III. The C-terminal (CT) dark blue. Sequence between the linker region and the beginning of the structure is indicated by a dotted line in light gray. **b** Phylogenetic tree of sequences of α/β hydrolase core domain and if present the CT domain only from various members within the hydrolase superfamily. Each clade is indicated by a colored box naming the protein family, their known or predicted enzymatic function, and known metal ions essential for their respective enzymatic functions. **c** crystal structure of YejM PD with indicated location of active site. **d** LtaS soluble domain with indicated location of active site. **e** EptA periplasmic domain with indicated location of active site.

Structures of two YejM PD homologues show similarity to hydrolase-fold enzymes like arylsulfatases and alkaline phosphatases ^26^. In spite of very low sequence similarity, the α/β hydrolase core structure of YejM PD is similar to members of the hydrolase superfamily, including the extracellular domain of lipoteichoic acid synthase (LtaS) of *Staphylococcus aureus* ^26^. The soluble domain structure of LtaS revealed a bound Mn^2+^ ion at the enzyme active site near the catalytic nucleophile threonine residue ^13^. Structures of PEA transferases like EptA and MCR show the same fold and an active site with bound metal ions and substrates. The conserved and crucial catalytic nucleophile threonine residue is especially important in catalysis since bacteria showed decrease inhibition by CAMPs, such as colistin and polymyxin, when substituted to alanine in MCR-1 ^27–29^.

How YejM facilitates translocation of cardiolipin is unknown. YejM forms tetramers *in vitro*, however, its physiological oligomeric state and whether it acts as a channel, transporter or in conjunction with another protein is unknown. Without known enzymatic activity or co-factors, it is unclear where the energy for cardiolipin translocation comes from. To date, no enzymatic activity has been reported for YejM ^22,26^.

Here, we report a novel enzymatic function of YejM. We have solved multiple high-resolution crystal structures of wildtype and mutant YejM PD that reveal an intact active site with bound metal ions and highly conserved catalytic nucleophile threonine residue. We demonstrate that YejM has a phosphatase activity that is dependent on the presence of magnesium ions. Further, we show that the enzymatic activity of YejM depends on the presence of its intact active site and the 5TM domain, and that increased levels of cardiolipin in the OM depend on enzymatic activity of YejM.

## Results

### Identification of the active site in YejM and structural homologues

We solved three crystal structures of the periplasmic domain of YejM (YejM residues 241-586) from *S. typhimurium*: two crystal structures of the wildtype YejM PD, and a third with a F349 to alanine mutation to the resolutions of 2.35 Å, 1.92 Å, and 2.05 Å, respectively (Table 1). Their general architecture resembles the α/β hydrolase fold, made up of alternating α-helices and β-sheets forming three layers (Figure 1a) ^26^. The α/β hydrolase core is the landmark domain shared within the large hydrolase superfamily including sulfatases, phospho-, mono-, and diesterases, and metalloenzymes (Figure 1b; Supplemental Table 1). Compared to other enzymes, YejM has an additional <u>C</u>-terminal (CT) domain of unknown function (Figure 1c). This CT domain shows structural similarity to a few other proteins such as colicin and kinases such as PLK1 (Supplemental Table 2). Interestingly, in some instances, this domain acts as a site for oligomerization ^30^.

**Table 1:** X-ray data collection and refinement statistics for YejM PD structures.

|  | **YejMPD** | **YejMPD-twinned** | **YejMPD-F349A** |
| --- | --- | --- | --- |
| PDB ID | 6VAT | 6VDF | 6VC7 |
| Data collection | ALS BEAMLINE 4.2.2 | ALS BEAMLINE 4.2.2 | ALS BEAMLINE 4.2.2 |
| Space group | P 32 2 1 | P 1 | P 21 21 21 |
| **Cell dimensions** |  |  |  |
| a, b, c (Å) | 113.80,113.80,299.78 | 42.948, 42.957, 181.209 | 121.45,125.10,182.75 |
| $\alpha$, $\beta$, $\gamma$ (°) | 90.00, 90.00, 120.00 | 94.28 94.015 111.779 | 90.00,90.00,90.00 |
| Resolution (Å) | 59.655-2.35 (2.39-2.35)* | 59.9-1.92 (1.989-1.92) | 47.50-2.05 (2.08-2.05) |
| Rmerge | 0.224 (2.192) | 0.08983 (0.7619) | 0.112(2.078) |
| I / σI | 11.1 (0.7) | 7.85 (1.07) | 13.8(0.6) |
| Completeness (%) | 99.9 (99.0) | 77.15 (60.59) | 85.88 (48.97) |
| Redundancy | 9.6 (5.0) | 1.6 (1.4) | 6.4 (3.7) |
| **Refinement** |  |  |  |
| Resolution (Å) | 56.9 - 2.35 (2.434 - 2.35) | 59.9 - 1.92 (1.989 - 1.92) | 43.57 - 2.05 (2.123 - 2.05) |
| No. reflections | 94485 (9231) | 70265 (5488) | 150131 (8464) |
| R<sub>work</sub> / R<sub>free</sub> | 0.1963/0.2489 | 0.2141/ 0.2639 | 0.1847/0.2270 |
| No. atoms | 17290 | 11745 | 17703 |
| Protein | 16358 | 10872 | 16353 |
| Ligand/ion | 245 | 61 | 340 |
| Water | 687 | 812 | 1010 |
| B-factors | 52.67 | 20.42 | 47.21 |
| Protein | 53.00 | 19.82 | 47.01 |
| Ligand/ion | 61.84 | 47.33 | 62.56 |
| Water | 41.64 | 26.37 | 45.18 |
| R.m.s deviations |  |  |  |
| Bond lengths (Å) | 0.004 | 0.002 | 0.005 |
| Bond angles (°) | 0.62 | 0.48 | 0.77 |
| **Ramachandran statistics** |  |  |  |
| favoured (%) | 92.11 | 90.47 | 94.59 |
| allowed (%) | 6.38 | 7.84 | 4.39 |
| outliers (%) | 1.51 | 1.69 | 1.02 |

\*Statistics for the highest-resolution shell are shown in parentheses.

Despite their high sequence variability, proteins from the hydrolase superfamily have a conserved metal-specific active site that is located at the base of layers II and III of the hydrolase domain (Figure 1 c, d, e). Most of these proteins are metalloproteins (Supplemental Tables 1 and 2). YejM has predicted structure and domain similarities with bacterial proteins involved in antibiotic resistance. This includes MCR proteins, ^14,29^, EptC from *Campylobacter jejuni*, EptA from *Neisseria meningitidis*, etc. (Supplemental Table 2). Unlike YejM and EptA/B/C, which are encoded on the chromosome, the genes encoding MCR1 proteins are located on plasmids and highly transferable across species ^14,31^.

We identified a metal ion binding site in YejM that is conserved across many members of the larger phosphatase super-family (Figure 1b), and is located at the base of layers II and III (Figure 1c and Figure 2a-c). Phylogenetic analysis of periplasmic domains only across selected members of the phosphatase super-family revealed that YejM is more closely related to the LtaS in Gram-positive bacteria, and more distantly to PEA transferases EptA and MCR-1 in Gram-negative bacteria (Figure 1b).

**Figure 2.**
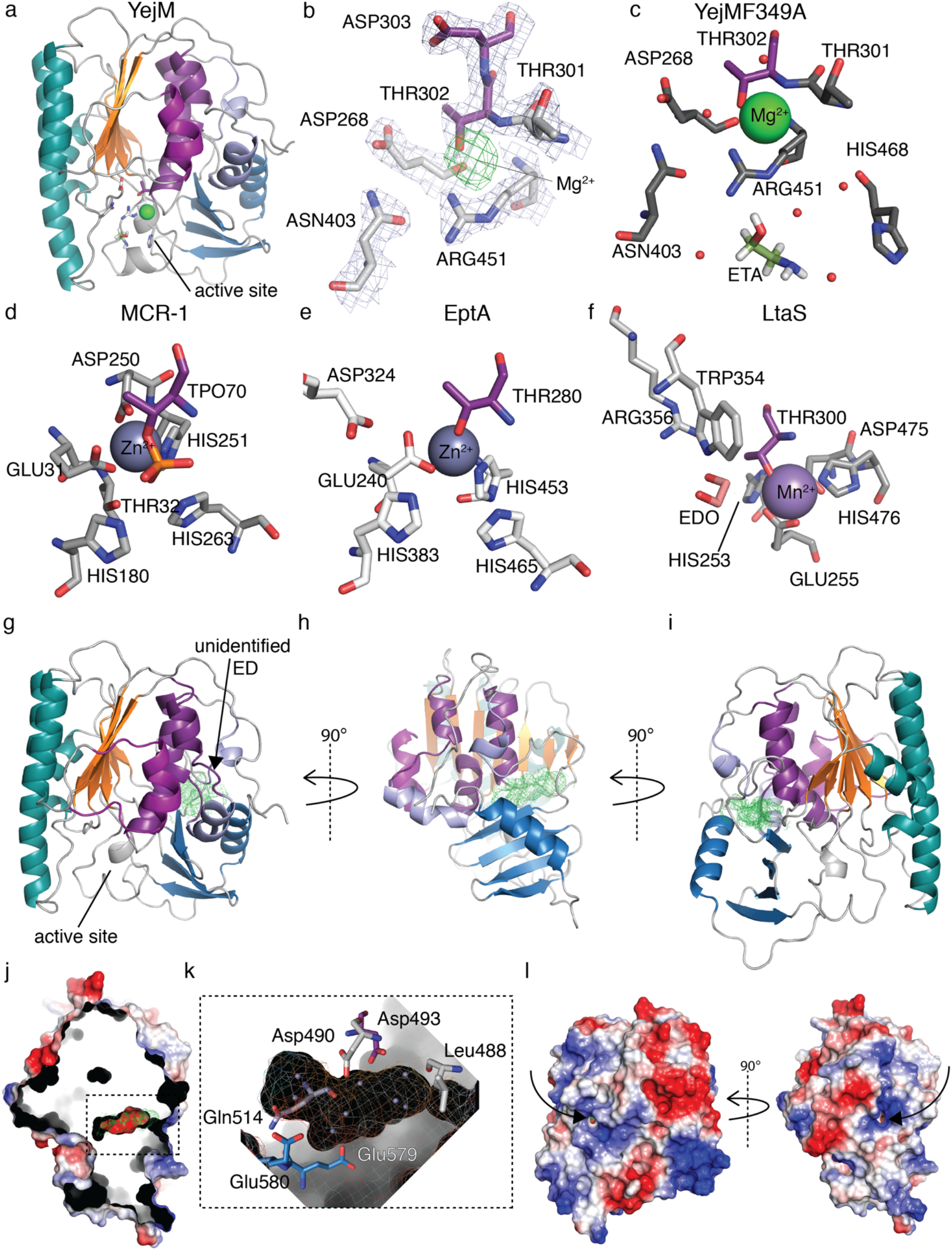
Location of novel YejM active site and substrate sites. **a** YejM PD with magnesium ion (green) in a larger vestibule at active site below layer II and III. **b** Active site of YejM PD F349A with conserved threonine Thr302 (deep purple) from layer III, magnesium (Mg^2+^) shown by 2Fo-Fc electron density map (green mesh) at sigma level 2.5, Thr301, Thr302, Asp303, Asp268, Asn403, and Arg451 with 2Fo-Fc electron density (light blue mesh) at sigma level 1.5. **c** Active site of YejM PD F349A with conserved threonine Thr302 (deep purple) from layer III, other residues involved in Mg^2+^ (green sphere) and substrate ETA are Asp268, Asn403, Arg451 and His468, prospective specific waters (red spheres). **d** Active site of MCR-1 around zinc (Zn^2+^, blue sphere) include conserved phosphorylated threonine TPO70, Glu31, Thr32, His180, Asp250, His251, His263. **e** Active site of EptA around Zn^2+^ include conserved threonine Thr280, Glu240, Asp324, His383, His453 and His465. **f** Active site of LtaS with conserved threonine Thr300 (deep purple) from layer III involved in coordination of manganese (Mn^2+^, purple sphere) and substrate EDO in stick representation orange, together with residues Glu255, Asp475, His253, Trp354, Arg356, His476. **g** Front view of YejM PD F349A, black arrow pointing towards the additional density observed in the Fo-Fc map at 2.2 sigma level (green mesh), located at interface between layer III and the CT domain. **h** YejM PD side view. **i** YejM PD back view. **j** Electrostatic surface presentation of YejM PD side view and cropped in Z plane to show large negatively charged pocket with additional Fo-Fc map (green mesh). **k** Zoom in on the negatively charged pocket lined by residues Leu288, Asp490, Asp493, Gln514 and Glu580. Dummy atoms (light blue spheres) placed into the pocket surrounded by the electrostatic surface overlaid in mesh and solid representation. **l** Electrostatic surface of YejM PD back and side views. Black arrows points towards positively charged access funnels (port 1) and (port 2) towards the visible Fo-Fc density (green mesh), indicating its accessibility from the outside.

### YejM has an active site similar to other metalloenzymes

All our YejM PD structures showed electron densities near the conserved threonine residue Thr302, which could not be satisfied with water molecules during structure building and refinement, but a Mg^2+^ or Mn^2+^ ion fit and refined well in it (Figures 2b,c). Multiple sequence and secondary-structure based alignments (Clustal Omega ^32^, Promals ^33^), revealed that the active site residues of LtaS and EptA are structurally aligned with those of YejM (Figure 2c-f). The metal coordination in all our YejM PD structures resembles metal coordination in MCR-1, EptA, and LtaS (Figure 2d-f). The active site coordination of YejM contains the conserved threonine 302 located at the base of layers II and III. Asp268, Asn403, Arg451 and His468 residues are involved in metal, water and substrate coordination (Figure 2b,c). We compared the active site coordination and found similarities to: i) MCR-1 with residues around its Zn^2+^ coordinated by Thr70 that is phosphorylated in this structure to TPO70, and further residues Glu31, Thr32, His180, Asp250, His251, His263; ii) EptA with residues around its Zn^2+^ ion coordinated by Thr280, and Glu240, Asp324, His383, His453 and His465 (Figure 2d,e). The active site of YejM resembles coordination of the active site around the Mn^2+^ of LtaS, with conserved threonine Thr300 and residues Glu255, Asp475, His253, Trp354, Arg356, His476 (Figure 2f). Although there are differences between the active sites, most residues and their charges, particularly the catalytically crucial nucleophilic threonine, are highly conserved across the active sites of other metalloenzymes (Supplemental Figure 1 and 2).

### The active site residues of YejM are highly conserved across homologues

To examine the active site residues conservation across YejM homologues, we chose three sequences centered around Asp268 (metal and substrate binding), Thr302 (catalytic residue and metal binding), and Arg451 (metal binding). In addition, we used secondary structure boundaries of the hydrolase domain to define these three sequences centered around Asp268, Thr302, and Arg451 (Supplemental Figure 2a). Our phylogenetic analysis revealed very high conservation for the sequence around Asp268 and Thr302, and more sequence flexibility around Arg451. Key residues Asp268, Thr302 and Arg451 show nearly 100% conservation across all homologues. Whereas Asp268 is 100% conserved across YejM homologues, Thr302 is substituted to an alanine in the clade including *Klebsiella pneumoniae* (KLEPN) and *Klebsiella michiganensis* (KLEMI) (Supplemental Figure 2a). Their sequence retained a threonine at the same place as YejM Thr301, which may compensate the loss of threonine at position Thr302. Two species from the clade comprised of *Enterobacter cloacae* (ENTCL) and *Raoultella terrigena* (RAOTE) indicate a serine substitution at Thr301; serine can also act as a nucleophile to form a covalent phospho-intermediate (Supplemental Figure 2a). At the position of Arg451 we observed a change to histidine in some species; this conserves the positive charge and characteristic of this site that is likely involved in metal binding (Supplemental Figure 2a). The major division between the clades are defined around conservation at His468. Whereas it is highly conserved amongst *E. coli*, *Salmonella* and *Citrobacter*, *Enterobacter* has an asparagine, and *Klebsiella* and *Raoultella* have a glutamine at this position (Supplemental Figure 2a,b). YejM homologues have slight differences around the conserved main active site residues and accommodate for various substrates and metal binding. Blasting our three initial *S. typhimurium* sequences (Supplemental Figure 2a) against the data base resulted in several conserved types of sequence alterations around all three active site sequences, building specific signature sequences that we found assembled across species (Supplemental Figure 1). Furthermore, we found that *E. coli* uses only type 1 and *S. typhimurium* only type 2 sequences, and other homologues used many combinations of various type of sequences (Supplemental Figure 1), which further increases the accommodation of substrate and metal use in a variety of enzymatic functions.

### YejM is a magnesium specific phosphatase

Based on conservation, location of the active site, presence of divalent metal ion, and the conserved threonine Thr302, we hypothesized that YejM has phosphatase activity. To test whether YejM can hydrolyze phosphate groups, we chose the fluorogenic phosphatase substrate 6,8-Difluoro-4-Methylumbelliferyl Phosphate (DiFMUP) as a substrate. To assess whether enzymatic activity is dependent on metal ions, we performed the assay with and without equimolar amounts of salts of various divalent cations. Our data showed for the first time that YejM has a phosphatase activity, which is highly specific to the presence of magnesium ions (Figure 3a). YejM in presence of magnesium (Mg^2+^) resulted in a 100-fold increase in the phosphatase activity compared to other metals or without additional metal ions or added EDTA (Figure 3a). Thus, YejM has a phosphatase activity that is specifically dependent on the presence of Mg^2+^.

**Figure 3.**
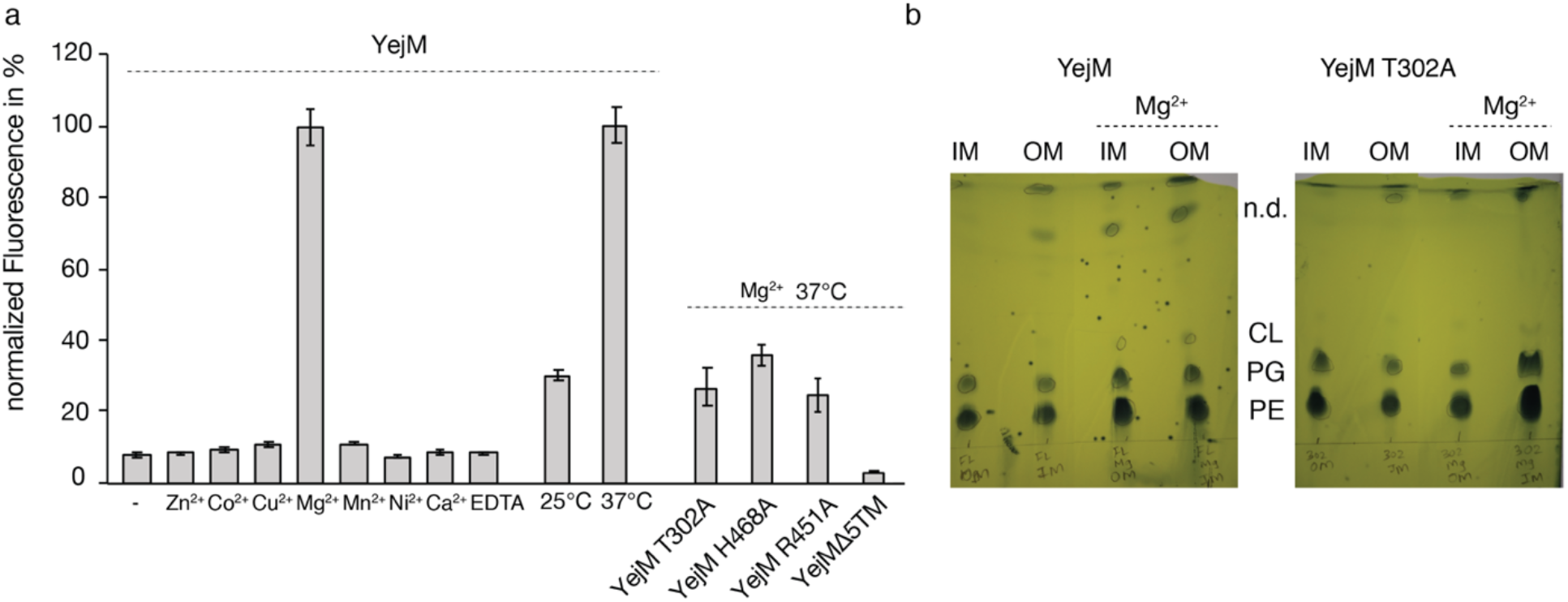
YejM is magnesium dependent phosphatase and cardiolipin translocation to the OM is coupled to Mg^2+^ enzymatic activity of YejM. **a** Phosphatase activity of YejM in the absence and presence of various divalent metals and EDTA. Equimolar amounts of divalent cations (Zn^2+^, Co^2+^, Ni^2+^, Mn^2+^, Cu^2+^, Mg^2+^, and Ca^2+^), and EDTA were added to the reaction. The maximum activity (in the presence of Mg^2+^) was set to 100 % and all other activities were scaled accordingly. Phosphatase activity of YejM at different temperatures (25 and 37 °C). Activity of YejM at 37 °C was set to 100 % and activity at 25 °C was scaled accordingly. Phosphatase activity of wild type YejM, YejMT302A, YejMH468A and YejMΔ5TM. Activity of wild type YejM was set to 100 % and remaining activities were scaled accordingly. All data are an average of three experiments. Error bars indicate the standard deviation of the data. **b** Inner membrane IM and outer membrane OM preparations of cells expressing YejM or YejM Thr302A (T302A) without and with additional magnesium (Mg2+) in the media. Lipids were detected by phosphomolybdic acid. Inner membrane (IM) and outer membrane (OM). Detected lipids are phosphatidylglycerol (PG), phosphatidylethanolamine (PE), cardiolipin (CL), and not determined lipid species (n.d.).

The observed enzymatic activity of YejM is lower compared to potato acid phosphatase (PAP) (Supplemental Figure 3), which may be attributed to the following two factors: i) DiFMUP might differ substantially from the natural substrate for YejM; ii) the absence of a second substrate serving as a phosphate acceptor molecule might slow down the substrate turnover rate of YejM. We tested whether the activity of YejM is temperature dependent and conducted the assay at 25°C and 37°C. Indeed, YejM has a 3.3-fold higher phosphatase activity at 37°C as compared to the activity at 25°C (Figure 3a). Higher activity at 37°C suggests that YejM increases activity upon host-infection, at which point Gram-negative bacteria are exposed to body temperatures higher than 36°C, and therefore YejM is most effective in OM modification for temperature adaptation and change in permeability.

We performed alanine substitution mutagenesis at active site residues and tested activity. Mutation of Thr302 to alanine resulted in about 70% loss of phosphatase activity (Figure 3a). Substitution of His468 to alanine, which resembles the similar structural location of His465 and His476 in EptA and LtaS, respectively, resulted in an approximately 60% decrease of phosphatase activity (Figure 3a). Next, we tested whether the periplasmic domain alone has enzymatic activity. We used YejMΔ5TM lacking the 5TM domain, which showed a drastic loss of ~97% phosphatase activity and is lower than YejM with divalent cations other than magnesium (Figure 3a). Our results suggest that both, the active site and the 5TM domain including the linker region are essential for the enzymatic activity of YejM.

### Increased cardiolipin levels in the OM are linked to the enzymatic activity of YejM

YejM is associated with increased amounts of cardiolipin in the OM and is proposed to be part of an essential cardiolipin transport system ^22,26^. Since we found an intact active site, bound metal ions and enzymatic activity, we asked the question whether the increase of cardiolipin into the OM is directly coupled to the magnesium specific enzymatic activity of YejM. To test this, we analyzed the lipid composition of the IM and OM of *E. coli* grown under various conditions and analyzed equal amounts of membranes for their lipid content using thin layer chromatography (TLC). We observed increased cardiolipin levels in the IM and OM from cells overexpressing YejM in growth media that was supplemented with 10mM MgCl_2_ (Figure 3b). In contrast, IM and OM samples from cells growing without magnesium supplement resulted in cardiolipin levels under the detection limit of TLC (Figure 3b). Additionally, an unidentified lipid species was detected in both membranes when YejM was expressed with increased amounts from cell membranes derived from magnesium supplemented media (Figure 3b).

These results show that cells growing in the presence of higher magnesium and overexpression of YejM present more cardiolipin in IM and OM. Next, we tested whether the increase in cardiolipin is dependent on the intact active site of YejM. Under the same growth conditions, with and without addition of magnesium, we overexpressed YejM carrying the Thr302A mutation that showed reduced activity (Figure 3a). IM and OM membranes presented cardiolipin levels that were nearly below the detection limit. However, the highest amount of cardiolipin could be detected in the OM of cells derived from magnesium supplemented growth media (Figure 3b). We conclude that increased levels of cardiolipin are linked to the magnesium-dependent enzymatic activity of YejM.

### YejM has a potential secondary substrate site between the hydrolase and C-terminal domains

Whether cardiolipin binds to the active site or is being translocated by the energy generated from catalysis of a yet unknown substrate that binds to the active site is not known. To dephosphorylate Thr302 YejM might require binding of a second substrate or an accessory protein. Following the hypothesis that a secondary substrate or protein may bind to the PD domain, we analyzed existing cavities in our structures (Figures 2d-l, Figures 4 and 5). A hydrophobic cavity located between layers II and III within the α/β hydrolase domain was proposed to bind the acyl chains of cardiolipin during translocation across the periplasmic space (Figure 4 and 5) ^26^. This potential cardiolipin binding pocket is lined by hydrophobic amino acids, namely, Val262, Phe292, Leu343, Phe394, Trp396, Val471, Leu473, Trp477, occurring on the β-sheets of layer II, and residues Leu309, Phe310, Leu334, Phe349, Tyr354, and Phe519 of layer III. Phe292 is absolutely essential for cell survival, and Phe275, Phe349, Phe362, and Trp396 were essential for growth in antibiotic-containing media ^26^. A proposed lid including residues 345-370 is hypothesized to act as a gate that closes or opens for acyl chain binding into this hydrophobic cavity ^26^ (Figure 5). The position of this loop in the six monomers in the asymmetric unit of our YejM PD structures differ by as much as ~8 Å, similar to the previously reported structures ^26^ (Supplemental Movie 1).

In addition to the hydrophobic cardiolipin binding cavity, we observed an unidentified electron density located in a negatively charged cavity at the interface between the hydrolase and the CT domain of in the YejM-PDF349A structure (Figure 2g-j). Contributing to its negatively charged electrostatic surface are residues Asp488, Asp490, Asp493, Gln514, Glu579, and Glu580 (Figure 2k). This electron density remained unidentified despite of our efforts using various methods including modeling and refining the structure with various possible ligands and identification by mass spectrometry. Interestingly, this negatively charged pocket is accessible by two mostly positively charged funnels in the structure (Figure 2l). The location, electrostatic nature and its accessibility makes it an intriguing site to bind a second substrate. We looked for similar negatively charged pockets at similar locations in other structures and identified one in LtaS (Figure 4a-f). No function is reported for this site for LtaS or other sulfatases and phosphatases. Our findings suggest that YejM may bind a second substrate that serves to dephosphorylate Thr302.

**Figure 4.**
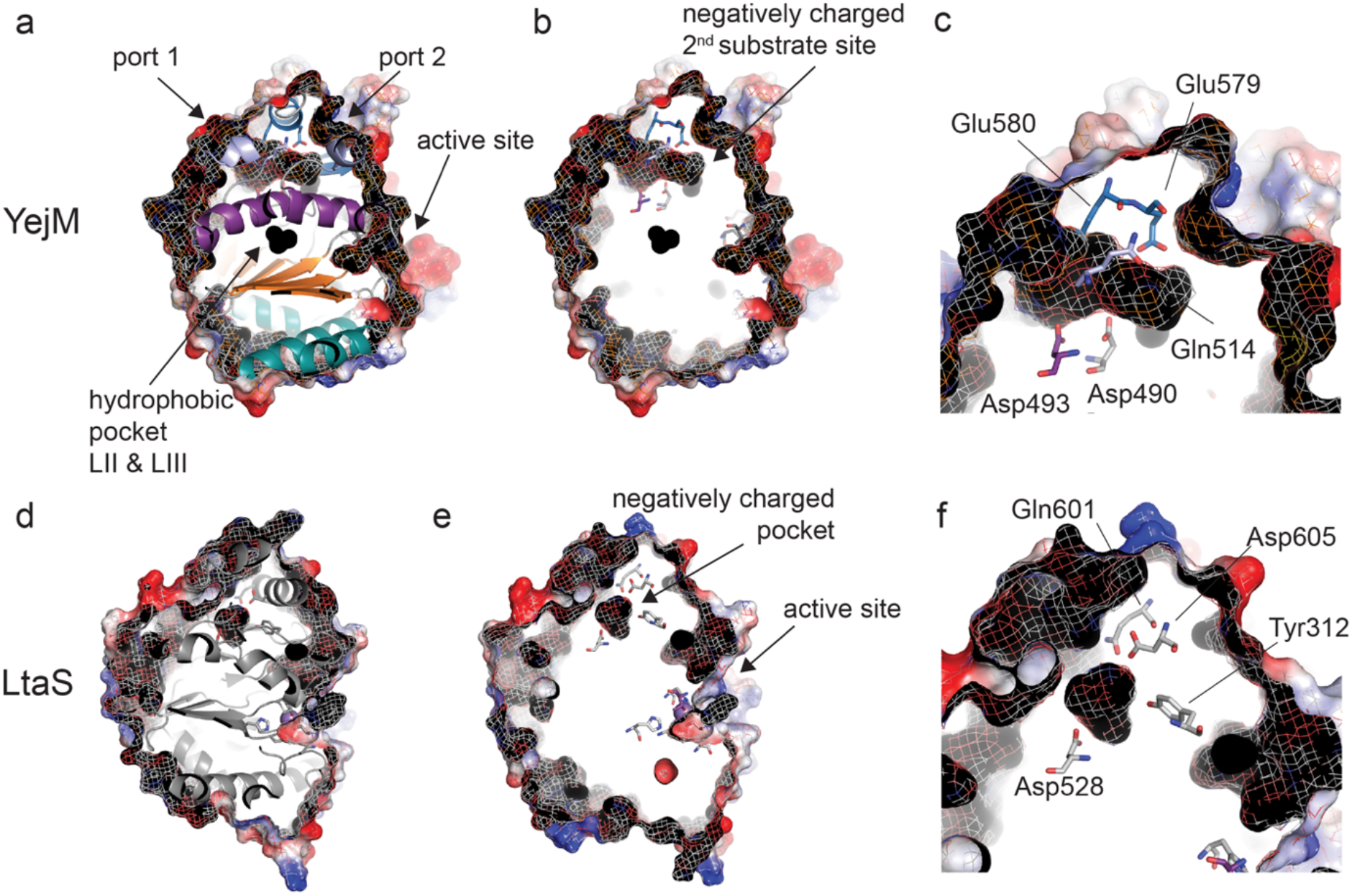
Secondary substrate pocket site of YejM in comparison to LtaS. **a** YejM PD viewed from the top showing cartoon representation of the peptide chain and electrostatic surface representation in an overlay of solid and mesh surface, structure is cropped in Z plane to show hydrophobic pocket (black arrow) and negatively charged secondary substrate side pocket and ports 1 and 2 (black arrows). Location of the active site is indicated by black arrow to the front of the PD domain. **b** YejM PD viewed from the top showing only electrostatic surface representation in an overlay of solid and mesh surface, structure is cropped in Z plane to show hydrophobic pocket and negatively charged secondary substrate side pocket (black arrow). Location of the active site is indicated by black arrow to the front of the PD domain. Residues located around the negatively charged pocket and towards the active site are shown in stick representation. **c** Zoom in to the location of the negatively charged secondary substrate side pocket with residues in stick representation. **d** LtaS viewed from the top showing cartoon representation of the peptide chain and electrostatic surface representation in an overlay of solid and mesh surface, structure is cropped in Z plane to show a pocket at a similar location to YejM PD negatively charged pocket at proposed secondary substrate site. **e** LtaS in same view as in D without cartoon representation of the main peptide chain. Residues lining a negatively charged pocket and the active site are shown in stick representation. **f** Zoom in to the location of the negatively charged pocket of LtaS with lining residues in stick representation.

**Figure 5.**
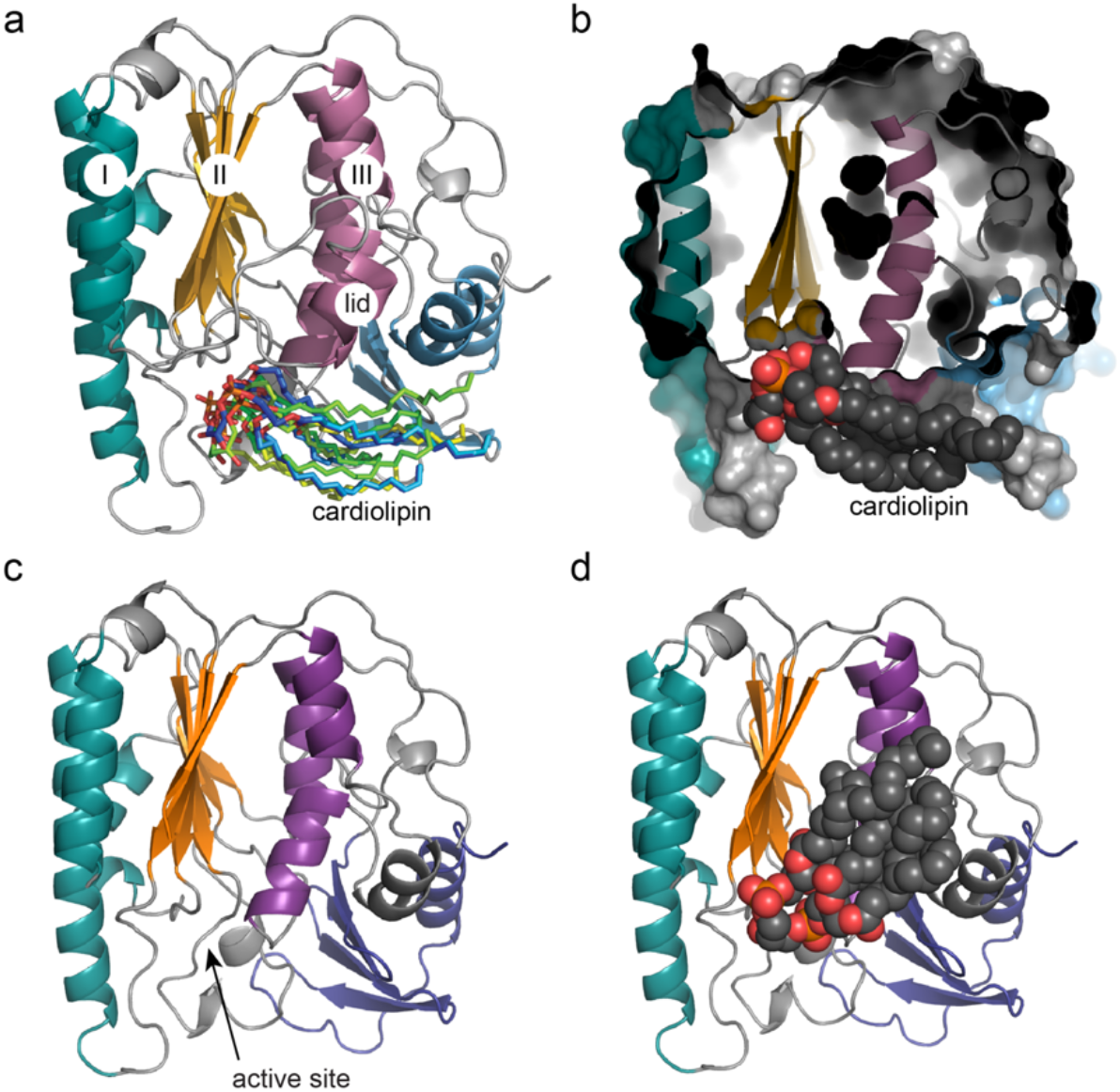
Cardiolipin docking to YejM PD with and without lid. **a** YejM PD structure indicating layer I, II and III, intact lid, and superimposed cardiolipin molecules in their preferred docked location and orientations, with phospholipid heads groups pointing towards the active site and acyl chains towards the CT domain. **b** YejM PD shown in cartoon and surface volume representation cropped in the Z-plane to visualize the vestibule of the active site, Black arrow point towards preferred location of cardiolipin molecule. Gray arrow points towards the hydrophobic pocket and proposed cardiolipin binding site between layers II and III. **c** YejM PD with removed lid, black arrow pointing towards the exposed area of the hydrophobic pocket between layers II and III. **d** YejM PD with removed lid shows cardiolipin molecule with preferred acyl chains flipped upwards towards the hydrophobic pocket while the phospholipid headgroups remain located towards the active site vestibule at the base of layers II and III.

## Discussion

We have unveiled a new functional identity of the essential inner membrane protein YejM that is associated with increased concentration of cardiolipin to the OM during host infection. The structure determination of three YejM PD domains revealed a metal ion containing active site that is located at the base of the hydrolase domain. Phylogenetic analysis of the PD domain sequences revealed that the active site is conserved across many members of the larger phosphatase super-family. Key active site residues were identified in YejM that serve as important catalytic residues and metal binder. Phylogenetic analysis and structure-based sequence determination around the active site revealed that the active site residues of YejM are highly conserved across homologues but also indicate changes that likely accommodate for different substrate and metal binding. Our enzymatic assay show that YejM has magnesium dependent phosphatase activity. The integrity of the active site and the 5TM domain including the linker region are essential for YejM enzymatic activity. Cardiolipin translocation by YejM is directly coupled to its enzymatic function. A second substrate binding site indicates that YejM might function in a multi-step process that combines enzymatic activity with cardiolipin translocation. Taken together, we assign a new functional identify to YejM: a metalloenzyme whose activity is coupled with translocation of cardiolipin to the OM. Our results bring intriguing questions to light about how YejM functions as an enzyme, the nature of its substrates and how it works on a molecular level, and how it is coupled to increased levels of cardiolipin in the OM. Our results fundamentally change how we think about this essential membrane protein that ultimately achieves survival of Gram-negative bacteria during infection.

Our extensive structure similarity search provided us with many enzymes to which we compared YejM with (Supplemental Tables 1 and 2). Despite medium to low overall sequence identity, the high sequence conservation at key active site residues allowed transfer of knowledge from better studied enzymes to begin to understand the newly identified enzymatic nature of YejM. YejM has previously been compared to an arylsulfatase and the lipoteichoic acid synthase LtaS ^26^. Despite their striking structural similarities, the authors concluded that because of lack of crucial cysteine and serine residues and the lack of metal ions in their structures, YejM cannot function as an arylsulfatase and is likely not a metalloenzyme ^26^. However, our structures, phylogenetic and biochemical data suggest that YejM is indeed a metalloenzyme with a conserved active site similar to LtaS, EptA and MCR1, and has phosphatase activity that is dependent on magnesium ions and the integrity of the active site (Figures 2b-f, 3a,b). Interestingly, the sequence of the YejM periplasmic domain is more closely related to LtaS in Gram-positive bacteria than pEtN transferases EptA and MCR-1 of Gram-negative bacteria (Figure 1b), suggesting YejM and LtaS may be distant orthologs. Members of the larger alkaline phosphatase family balance specificity and promiscuity in their evolution around the active site resulting in multidimensional activity transitions that may also hold true for the sub-family YejM belongs to ^34–36^.

Our data show that in the presence of magnesium, YejM can remove the phosphate group from DiFMUP (Figure 3a). We speculate that the Mg^2+^-specific enzymatic activity of YejM was missed in previous studies. This function would place YejM to be enzymatically closer to EptA and MCR-1 than LtaS. EptA and MCR-1 both hydrolyze the phosphoethanolamine (pEtN) from phosphatidylethanolamine (PE), to then transfer the pEtN group to lipid A, which neutralizes lipid A and renders binding of polymyxin-group antibiotics ineffective. EptA transfers pEtN to lipid A that is presented by another protein and may bind two differently sized substrates at its active site ^12^. MCR-1/2 however are suggested to have a secondary substrate binding site and pEtN transfer to lipid A is facilitated by MCR proteins alone ^29,37,38^. The latter may be the nature of the negatively charged cavity in YejM (Figure 2g-l). We hypothesize that this negatively charged cavity serves as a secondary binding site for a substrate that acts as phosphate receptor to dephosphorylate Thr302. Interestingly, in our comprehensive *in silico* docking study and analysis based on the study by Planas-Iglesias ^39^, most negatively charged phospholipid head groups of cardiolipin localize towards the active site (Figure 5a-d). The acyl chain location flip towards the hydrophobic pocket between layers II and III when using a YejM PD model with missing lid sequence (Figure 5c,d). Our results suggest that YejM could bind two substrates and is involved in a multi-step process that combines enzymatic activity with cardiolipin translocation. We will need to identify the nature of both substrates and design specific assays to determine the true enzymatic nature and describe the catalytic mechanism of YejM. A full-length YejM structure will be necessary to reveal the ideal environment of the active site to bind metal ion and substrates including cardiolipin. We created a chimera model that that includes the 5TM and RR linker domains based on the full-length crystal structure of EptA (PDB ID 5FGN), and combined it with the PD domain of our YejM PD structure (Figure 6a,b). This model visualizes the possible location of the active site relative to the membrane plane, the positively charged linker region and loops and helices of the 5TM domain (Figure 6a,b). YejM couples cardiolipin translocation with the newly identified magnesium dependent enzymatic function. Interestingly, we observed proteolytic cleavage products for both YejM and YejM191-586 that suggest a cleavage site within the PD.

**Figure 6.**
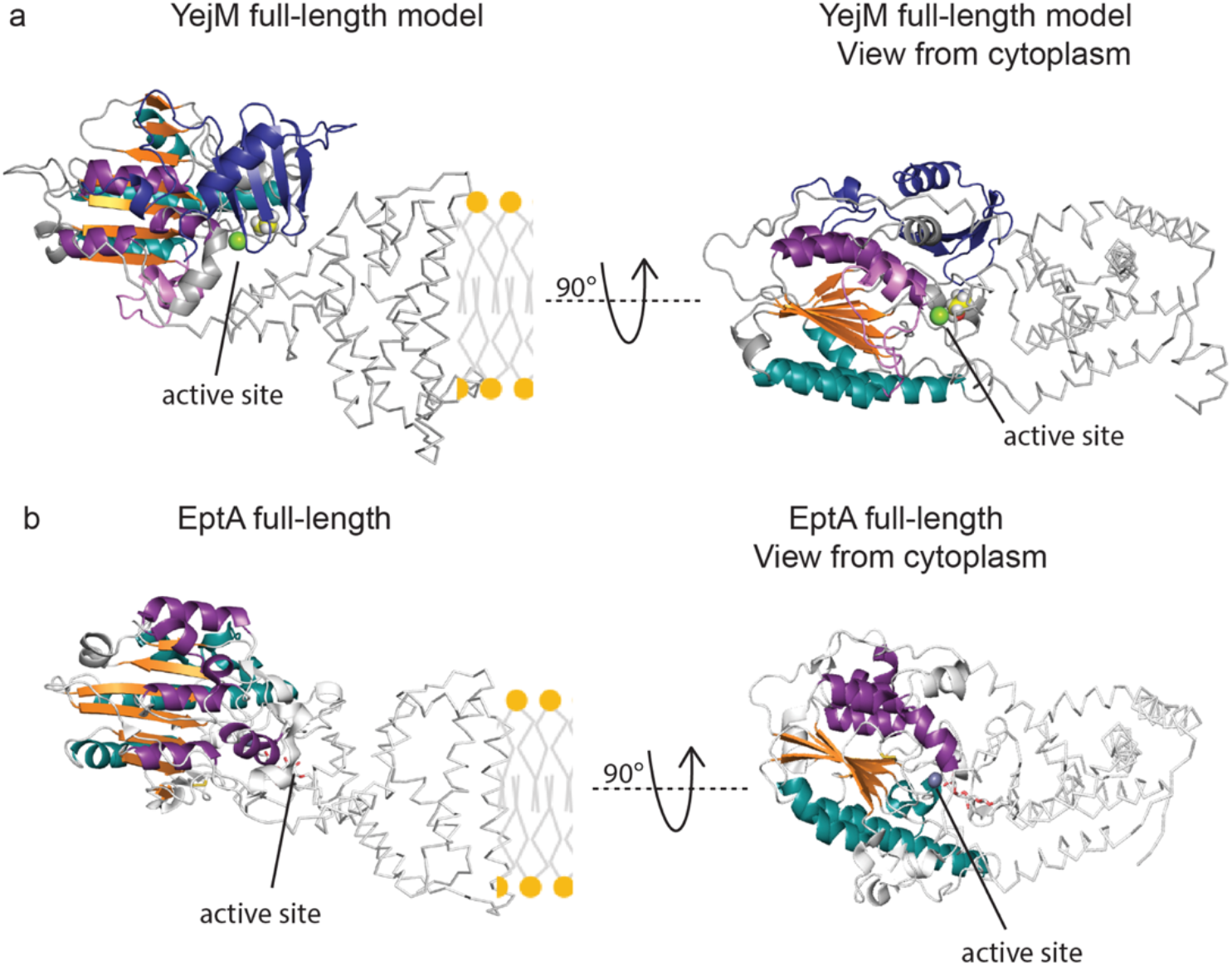
Full-length model of YejM. The YejM full-length model is a chimera between the model of the 5TM and RR linker domains based on the full-length crystal structure of EptA (PDB ID 5FGN) and the YejM PD from our original structure. **a** Left, side view of YejM full-length model within the membrane plane. Right, YejM full-length model rotated by 90° to view on top of the membrane plane from the cytoplasmic side. Location of the active site is indicated by line. **b** Left, side view of full-length EptA (PDB ID 5FGN) within the membrane plane. Right, EptA rotated by 90° to view on top of the membrane plane from the cytoplasmic side. Location of the active site is indicated by black line.

Together with our results, we speculate whether they have different functions that can work in concert with each other that are regulated by proteolytic processing. Notably, proteolytic cleavage has been observed in LtaS, which renders it inactive and controls its activity ^40^. Interestingly, a similar enzyme in *E. coli*, phosphoglycerol transferase OpgB, undergoes similar proteolytic cleavage that changes its enzymatic activity from phosphoglycerol transferase type I to type II ^41^. No proteolytic processing has been reported for EptA, MCR-1, and phosphoethanolamine transferase OpgE ^12,42,43^. We suggest proteolytic processing of YejM that separates 5TM from PD domain allows these two domains to interact with other proteins, separately.

While YejM might undergo proteolytic processing, both, the phosphatase and cardiolipin translocation activity depend on the integrity of the protein.

Specifically, the PD domain alone does not have enzymatic activity (Figure 3a).This is in stark contrast to EptA, for which the periplasmic domain alone was shown to retain full phosphatase activity ^27^. Similarly, the extracellular domain of LtaS alone is also fully active towards its substrate, phosphatidylglycerol ^44^. Notably, although the soluble domains of both EptA and LtaS are able to hydrolyze their substrates, EptA cannot add phosphoethanolamine to lipid A ^27^, and LtaS cannot synthesize lipoteichoic acid from glycerol phosphate monomers ^40^ in the absence of their transmembrane domains.

Interestingly, while the YejM 5TM domain alone is sufficient and essential for cell survival ^21,22^, it cannot translocate cardiolipin to the OM ^22^. Addition of the PD domain *in trans* to cells expressing only the 5TM domain does not rescue the loss of cardiolipin translocation function of YejM ^22^. This supports our results that cardiolipin transport to the OM is linked to the active site within the PD domain. Notably, Gram-negative bacterial strains without all three cardiolipin synthases (cls A, B, and C, *Δ*clsABC) can survive, whereas YejM deletion strain cannot ^45,46^. This suggests that cardiolipin translocation function of YejM alone is not the essential function for cell survival, even though it is important in pathogenicity of *S. typhimurium* and *Shigella flexnerii* ^22,45^. Interestingly, higher concentration of PG lipids were observed in the OM of *Δ*clsABC strain ^45^. Since YejM can bind PG as well as, we hypothesize that in the absence of cardiolipin, it might bind and translocate PG to the OM.

According to a recent study, YejM plays a phoPQ-dependent role in lipopolysaccharide assembly, that requires presence of an intact YejM. Absence of YejM PD leads to accumulation of lipid A-core molecules lacking O-antigen in the OM, as well as a decrease in amount of cyclopropanated fatty acids in the OM ^23^. These observations further strengthen the hypothesis that YejM influences LPS assembly and is involved in lipid homeostasis of bacterial membrane. It would be interesting to see whether these functions are further related to the enzymatic activity of YejM. Thus, based on the current body of research, YejM may have complex regulatory role apart from being a cardiolipin transporter that also depends on its enzymatic function.

Our study provides first important data giving a new functional identity to YejM that extends our understanding and provides a strong basis for future structure-guided approach to find inhibitors specific to YejM to fight infectious diseases.

## Material and Methods

### Mutagenesis

Site-directed mutagenesis were carried out to generate following mutants: YejMPD-F349A using forward (FW) and reverse (REV) primers: FW CTGTTTTCTTCGGATGGCGCCGCCAGCCCGCTTTATCGTC, REV GACGATAAAGCGGGCTGGCGGCGCCATCCGAAGAAAACAG; YejM-T302A using FW CATATGAGTTCAGGGAATACCGCTGATAACGGTATTTTCGGC, REV GCCGAAAATACCGTTATCAGCGGTATTCCCTGAACTCATATG; YejM-R451A using FW GTCGTGATCATTACCGCAGGAGCCGGCATACCGTTGACGCCG, REV CGGCGTCAACGGTATGCCGGCTCCTGCGGTAATGATCACGAC; and YejM-H468A using FW GTCGCAAGGTGCTCTGCAAGTAC and REV CAGTCGAAGCGATTTTCTTC. The mutagenesis sites are underlined. PfuTurbo DNA polymerase (Agilent Technologies) and Q5 Hotstart Mutatagenesis kit (NEB) were used.

### Expression and purification of YejM constructs

Full-length YejM cloned in pBAD24 vector and its mutants were expressed in *E. coli* Top10 cells and purified as described in ^22^. YejM-PD and its mutants, cloned in pET28 vector, were expressed in *E. coli* BL21(DE3) cells and purified described in ^47^

### Crystallization, X-ray diffraction data collection and processing

Purified YejM-PD wild type and mutant proteins were crystallized according to ^47^. Crystals were harvested using Litho loops (Molecular Dimensions) or Nylon loops (Hampton Research), and frozen in liquid nitrogen without or with paraffin as cryo-protectant. Diffraction data were collected at the Advanced Light Source beamline 4.2.2 in Berkeley, CA at 100K, using an oscillation of 0.1-0.2° per image. Diffraction images were indexed and integrated using XDS ^48^ and scaled and merged with Aimless using the CCP4 program suite ^49^.

### Structure determination

The structure of YejM241-586 from data set YejM-PD was solved in space group *P*3_2_21 by molecular replacement using Phaser with a monomer from the crystal structure of *Salmonella* PbgA globular domain 191 (PDB code 5I5F) as a search model ^26 50^. The structure was further refined using phenix.refine ^51^. Non-crystallographic symmetry (NCS) restraints were imposed in the early stages of refinement and released and instead Translation-liberation-screw motion (TLS) were applied during later stages of refinement. The structure was refined to a resolution of 2.35 Å. Data of YejM-PD-twinned and YejM-PDF349A were determined by molecular replacement using a monomer of YejM-PD (PDB ID 6VAT) as search model. Refinement of YejM-PD (YejM-PD-twinned) was carried to a resolution of 1.92 Å with applied detwinning operator *(-h,-k,h+k+l*) but otherwise following similar TLS refinement strategies as for YejM-PD. The structure of YejMPDF349A mutant was refined to 2.05 Å. The refinement statistics for all three structures are given in Table 1. Structures were deposited to the PDB database (https://www.rcsb.org/) and the following PDB IDs were assigned: 6VAT, 6VDF, and 6VC7.

### Phylogenetic analysis

The starting point of the phylogenetic analysis of YejM-related proteins were the *Salmonella thyphymurium* cardiolipin transport protein YejM (aka PbgA) sequence segments of interest:

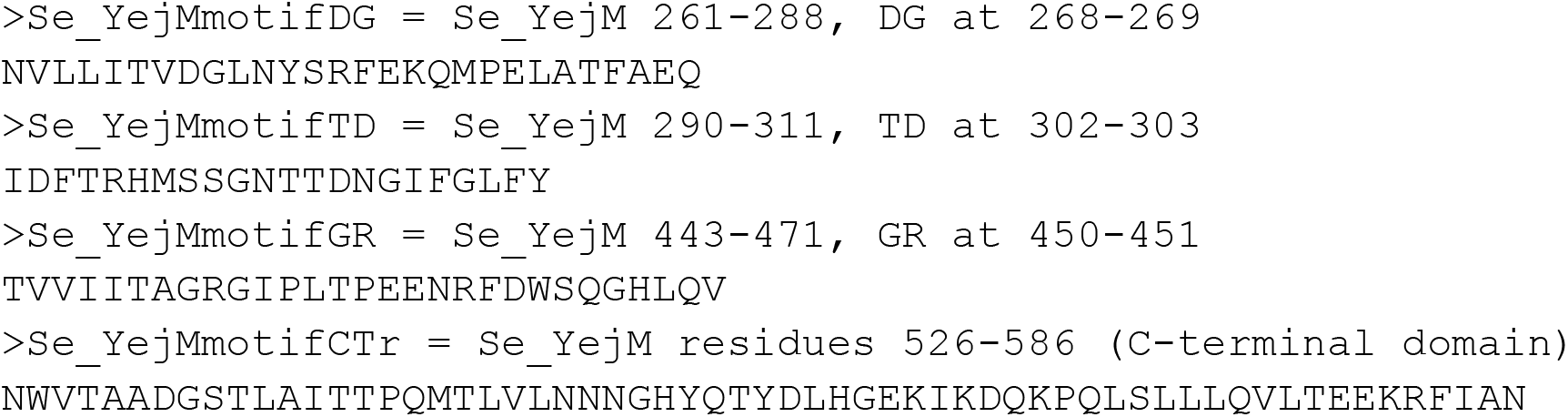

The first three segments contain conserved active site residues DG, TD, and GR motifs, respectively, while the last segments constitute the C-terminal domain of unknown function. Each of these sequences were used as input to *jackhmmer* of the HMMER v3.2.1 package (hmmer.org) in a search against the UniProt TrEMBL database (release 2019_04; locally installed) to derive profile hidden Markov models. With default parameters, a total of 2,933 sequences were identified in TrEMBL with significant matches to at least one of the motifs.

The alignment output of *jackhmmer* was processed to identify unique sequences. For example, there were 66 distinct sequences matching the DG motif and occurring at least four times. The input Se_YejMmotifDG occurred in 165 of the 2,933 TrEMBL sequences. These numbers reflect both redundancy of sequencing efforts (e.g., multiple recoveries of the entire YejM sequence from different samples) as well as evolutionary conservation across distinct organisms.

### Enzymatic assays

A 10 mM stock solution for DiFMUP substrate was prepared in N,N-dimethylformamide (DMF), stored at −20 °C, and used within 1 month. This stock was diluted to 200 μM to 1 mM in the respective reaction buffer just before use. For measuring the activity of YejM-FL, YejM-FL-T302A and YejM-FL-H468A, the reaction buffer consisted of 25 mM tris, pH 7.5, 150 mM NaCl, and 0.015 % DDM. For the periplasmic domain YejMΔ5TM, the reaction buffer used was 50 mM tris, pH 8.0, and 150 mM NaCl. In a typical experiment, 20 μM protein was incubated with 100 μM DiFMUP for minimum of four hours in dark. The temperature of incubation varied between 25 and 37 °C according to the experiment. Metal ion screen were conducted in the presence of equimolar amounts of salts of various divalent cations (ZnCl_2_, CoCl_2_, NiSO_4_, MnCl_2_, CuCl_2_, MgCl_2_, and CaCl_2_), or EDTA. Once the importance of Mg^2+^ ions in the phosphatase activity of YejM was confirmed, MgCl_2_ was included in 20X molar excess as compared to the protein in all subsequent assays. Appropriate negative controls (reaction buffers without protein and with same amount of MgCl_2_ and DiFMUP) were used for each experiment. Potato acid phosphatase (PAP) included in the assay kit was used as a positive control, at a working concentration of 0.1-1 U/ml. The reaction buffer for PAP was 50 mM sodium acetate, pH 5.0. All reactions were set in 100 μL volume, in 96-well black, flat bottom microplates (Greiner Bio-one, cat. no. 655209) covered with lids. The fluorescence of the product of the enzymatic reaction, 6,8-difluoro-4-methylumbelliferone (DiFMU), was measured using Synergy Neo2 plate reader (BioTek) by excitation at 358 nm and emission at 455 nm.

All experiments were conducted in triplicates and the average values were reported. Results from different experiments were compared by setting the final fluorescence value of the sample with maximum fluorescence to 100% and adjusting fluorescence values of other samples accordingly.

### Membrane lipid isolation

Total membranes were isolated from 1 L cultures of different strains of *E. coli* as per the protocol described in ^22^. Separation of inner and outer membrane fractions were performed as described by 45. Each membrane pellet was resuspended in 10 ml ddH_2_O and ultra-centrifuged at 50000 rpm for 45 min at 12 °C. After removing the supernatant, each membrane pellet was resuspended in 1 ml ddH_2_O. To this, 100 μL of 5% sarkosyl was added and incubated for 20 min at room temperature with shaking. Ultracentrifugation at 50000 rpm for 45 min at 12 °C yielded the inner membrane fraction in the supernatant. The pellet, that represents the OM fraction, was washed with 10 ml of 1% sakosyl and resuspended in 2 ml ddH_2_O.

Lipids were isolated from the IM and OM fractions by the method of Bligh and Dyer ^52^. Briefly, to each 1 ml of sample in a glass vial, 3.75 ml 1:2 (v/v) chloroform:methanol was added and mixed by vortexing. This single-phase Bligh-Dyer layer was converted into a double phase by adding 1.25 ml Chloroform followed by 1.25 ml ddH_2_O and vortexing. The two layers were separated by centrifuging the glass vials at 2000 RPM in a table-top centrifuge for 10 min at room temperature. The bottom phase was recovered by carefully aspirating with a glass pasture pipette and transferring to another glass vial. The original aqueous phase was washed with another 1.25 ml of Chloroform and vortexing and centrifuging as before. The organic phase from this wash was mixed with the previous one. The lipids were dried under vacuum and resuspended in such a volume of 4:1 chloroform:methanol as to obtain 1 mg/ml final concentration of lipids.

### Thin Layer Chromatography

Thin-layer chromatography (TLC) was performed by using Silica Gel 60 Aluminum Sheets (MilliporeSigma™; catalog no. M1055530001). 10 cm X 10 cm squares were equilibrated with 1:1 chloroform:methanol in a Fungicrom separating TLC chamber (Fungilab™; catalog no. 06-815-188). After air drying the plates in a fume hood for an hour, the lipids were spotted (5-15 uL of 1 mg/ml of membrane solution) at two cm from the bottom of the plate and 2 cm apart from each other. The plates were air dried completely. The lipids in the membrane samples were then separated in the TLC chamber equilibrated in 50:30:10:5 chloroform:hexane:methanol:acetic acid until the solvent front reached 1 cm below the top. The plates were air dried and then stained with either iodine vapors, 20 % w/v phosphomolybdic acid (PMA) solution in ethanol (Santa Cruz Biotechnology, catalog no. sc-203346), or 1:1 diluted molybdenum blue spray reagent (Millipore Sigma, catalog no. M1942-100ML). Developing spots on plates sprayed with PMA required heating with a blow dryer.

### *In silico* binding studies

The ideal coordinates of cardiolipin (CDL) were downloaded from the HIC-Up server ^53^ as a PDB file. AutoDock Tools (ADT) (version 1.5.6) ^54^ was used to add polar hydrogens to the protein and ligand molecules, calculate Gasteiger charges, and finally generating PDBQT files. ADT was also used to determine the dimensions of a grid box encompassing the entire protein chain. Actual docking was done using AutoDock Vina ^55^ using default docking parameters. Autodock Vina generates the output of docked ligand (9 conformations) in a PDBQT file, which was later converted in a PDB file and visualized using UCSF Chimera ^56^ and Pymol (The PyMOL Molecular Graphics System, Version 2.0 Schrödinger, LLC). The contacts between receptor and ligand molecules in various docked poses were determined using Contact program in CCP4 suite ^49^.

## Acknowledgements

We thank Dr. Jay Nix for his assistance at the ALS beamline 4.2.2 (http://mbc-als.org/), Dr. Giovanni Gonzalez-Gutierrez for his assistance in core-facilities at Indiana University Bloomington (Indiana University). We thank Dr. Volker Brendel (Indiana University) for help with phylogenetic analysis. Dr. Charles Dann III (Indiana University) for the use of BioTek Synergy Neo2 plate reader. We thank the following undergraduate students from the Ressl lab for technical assistance and discussions: Elaina Roach, Gene Qian, Rohit Das. We thank Dr. Julia van Kessel, Dr. David Giedroc, and Dr. Steven Bell (Indiana University) for discussions and critical comments on the manuscript. This project was funded through the Indiana University College of Arts and Sciences Bloomington startup funds to SR.

## Author contributions

U.G. designed research, performed experiments, analyzed data, and wrote manuscript. P.A.P.P. performed specific experiments, analyzed data, and revised manuscript. H.K. performed specific experiments. W.C. performed experiments. S.R. designed and directed research, performed specific experiments, analyzed data, prepared figures and wrote manuscript. All authors reviewed the manuscript.

## Data availability

Atomic coordinates and structure factors are deposited at the Protein Data Bank under the access codes 6VAT, 6VC7, 6VDF. Clones used in this study are available upon request to the corresponding author.

