## Supplemental Tables and Figures for "New functional identity of the essential inner membrane protein YejM: the cardiolipin translocator is also a metalloenzyme"

**Title**

**Authors Affiliations**

**Supplemental material**

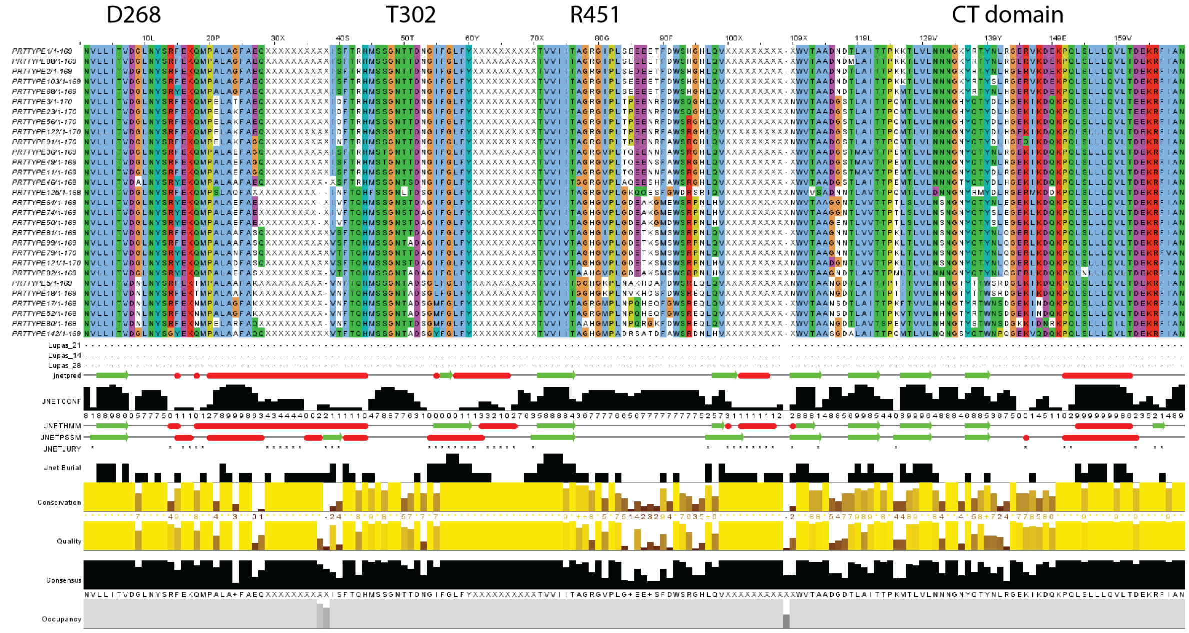


**Supplemental Figure 1 Sequences around the active site residues and CT-domain, their conservation and natural occurring variations.** The first stretch of sequence is centered around aspartate D268, the second around threonine T302, the third around arginine R451, and the fourth covers the CT domain.

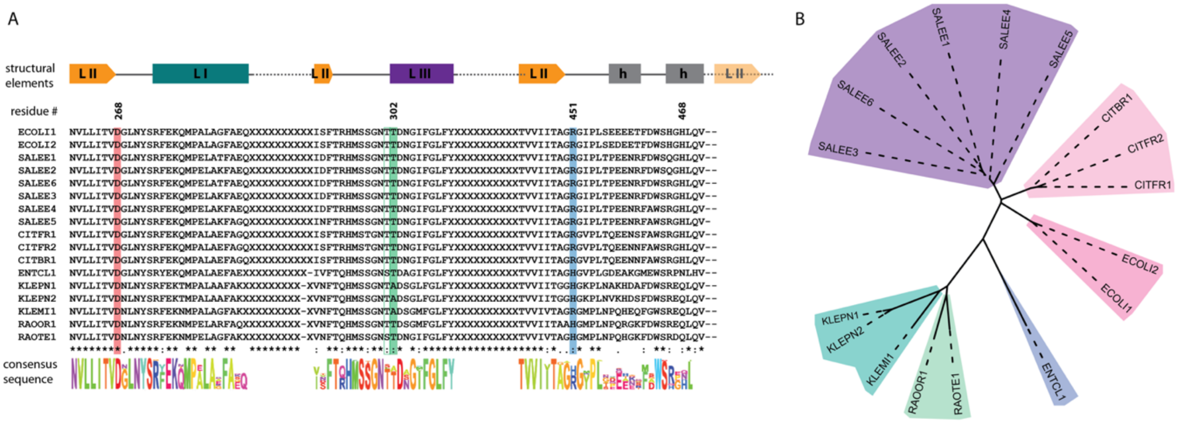


**Supplemental Figure 2 Active site residue conversation across YejM homologues. A** Sequence alignment of all three sequences around residue Asp268 (underlaid with red box), Thr302 (underlaid with green box), Arg451 (underlaid with blue box). Structural elements that guided our choice of sequence boundaries around those sites are indicated on the top representing beta-sheets with right facing rectangular arrow, helices with rectangles, and loops elements with a single line. Spaces between the sequence elements are marked with a dotted line. The color of the secondary elements reflects the same color code for layers I, II and III as in Figure 1. Layer I turquoise, layer II orange and layer III deep purple. The consensus sequence is shown at the bottom of the sequence, the color scheme is random and for better readability only. **B** Phylogenetic tree of YejM homologues. The length of branch lengths reflects distances between the clades. The unrooted tree groups YejM homologues of *Klebsiella pneumoniae* (KLEPN), *Klebsiella michiganensis* (KLEMI), *Raoultella terrigena* (RAOTE) and *Raoultella ornithinolytica* (RAOOR) into one clade (blue-green), *Enterobacter cloacae* (ENTCL) builds a separate outgroup (blue), various strains of *Salmonella* (SALEE1-5) together with *Escherichia coli* (ECOLI) and *Citrobacter braakii* and *freundii* (CITBR, CITFR, respectively), build a large clade (pink to purple). Phylogenetic tree was created with iTol (itol.embl.de) [57].
.

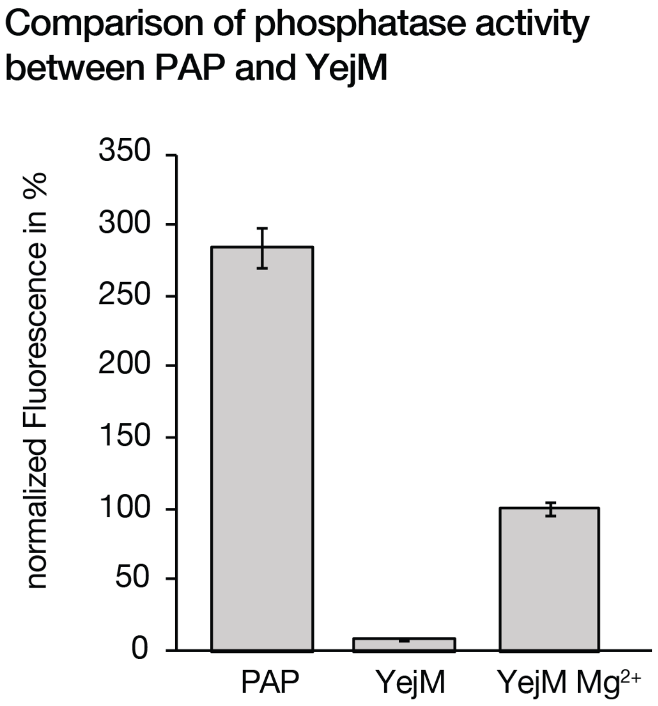


**Supplemental Figure 3** YejM enzymatic activity for substrate DiFMUP in comparison to potato acid phosphatase (PAP). YejM enzymatic activity is presented in normalized fluorescence units (y axis). YejM with additional $Mg^{2+}$ has a 1/3 o*f* PAP enzymatic activity for this substrate and YejM without addition of $Mg^{2+}$ has close to no observable activity.

**Supplemental Table 1: DALI comparison of YejM PD to other hydrolases/isomerases/transferases**

| PDB | Protein name | Enzyme family | Organism | Z score | rmsd | lali | nres | %id | Metal ion |
|---|---|---|---|---|---|---|---|---|---|
| 6j66 | Chondroitin sulfate/dermatan sulfate endolytic 4-O-sulfatase | Hydrolase | *Vibrio sp.* FC509 | 30.7 | 2.7 | 308 | 476 | 17 | $Ca^{2+}$ |
| 3b5q | putative sulfatase | Hydrolase | *Bacteroides thetaiotaomicron* | 30.5 | 2.6 | 305 | 467 | 17 | $Zn^{2+}$ |
| 4miv | Sulfamidase | Hydrolase | *Human* | 30.3 | 2.8 | 305 | 480 | 18 | $Ca^{2+}$ |
| 3lxq | Uncharacterized protein | -- | *Vibrio parahaemolyticus* serotype O3:K6 (strain RIMD 2210633) | 30.2 | 3.4 | 316 | 409 | 14 | $Cl^{-}$ |
| 6hr5 | S1_25 family sulfatase module of the rhamnosidase FA22250 | Hydrolase | *Formosa agariphila* | 29.9 | 2.9 | 304 | 376 | 16 | $Ca^{2+}$ |
| 2w5q | LtaS | transferase | *Staphylococcus aureus* | 29.7 | 3.3 | 304 | 424 | 17 | $Mn^{2+}$ |
| 5g2v | N-ACETYL GLUCOSAMINE-6-SULFATASE | Hydrolase | *Bacteroides thetaiotaomicron* | 29.1 | 2.7 | 306 | 494 | 17 | $Ca^{2+}$ |
| 4upl | sulfatase SpAS2 | Hydrolase | *Silicibacter pomeroyi* | 28.2 | 2.7 | 307 | 555 | 14 | $Zn^{2+}$ |
| 5fql | iduronate-2-sulfatase | Hydrolase | Human | 28.1 | 2.9 | 302 | 507 | 19 | $Ca^{2+}$ |
| 1p49 | Placental Estrone/DHEA Sulfatase | Hydrolase | Human | 27.6 | 3.1 | 304 | 549 | 15 | $Ca^{2+}$ |
| 4cys | G6 mutant of PAS, arylsulfatase | Hydrolase | *Pseudomonas Aeruginosa* | 24.4 | 3.5 | 309 | 534 | 17 | Ca2+ |
| 5olt | extramembrane domain of the cellulose biosynthetic protein BcsG | Transferase | *Salmonella typhimurium* | 22.1 | 3.5 | 275 | 383 | 14 | $Zn^{2+}$ |

| | | | | | | | | | |
|---|---|---|---|---|---|---|---|---|---|
| 6c01 | ectonucleotide pyrophosphatase / phosphodiesterase 3 | Hydrolase | human | 20.1 | 2.9 | 247 | 819 | 13 | $Zn^{2+}$, $Na^{+}$, $Ca^{2+}$ |
| 5xwk | alkaline phosphatase | hydrolase | *Sphingomonas* | 20.1 | 2.8 | 239 | 530 | 15 | $Zn^{2+}$, $Ca^{2+}$ |
| 4b56 | ectonucleotide pyrophosphatase-phosphodiesterase-1 (NPP1) | Hydrolase | *Mus musculus* | 19.3 | 2.9 | 247 | 816 | 11 | $Zn^{2+}$, $Ca^{2+}$ |
| 6dgm | phosphoglycerol transferase GacH | Transferase | *Streptococcus pyogenes* | 18.5 | 4.0 | 266 | 382 | 12 | $Mn^{2+}$, $Ca^{2+}$ |
| 3igy | phosphoglycerate mutase | Isomerase | *Leishmania mexicana* | 17.5 | 3.1 | 228 | 549 | 16 | $Na^{+}$, $Co^{2+}$ |
| 5kxa | Ectonucleotide pyrophosphatase/ phosphodiesterase | HYDROLASE/ HYDROLASE INHIBITOR | human | 17.2 | 2.9 | 237 | 751 | 13 | $Zn^{2+}$, $Ca^{2+}$ |
| 4n7t | phosphopentomutase | ISOMERASE | *Streptococcus mutans* | 17.0 | 3.0 | 217 | 402 | 15 | $Mn^{2+}$ |
| 5m0s | Ectonucleotide pyrophosphatase/phosphodiesterase | HYDROLASE | *Rattus norvegicus* | 16.9 | 2.9 | 236 | 777 | 12 | $Zn^{2+}$, $Na^{+}$, $Ca^{2+}$ |
| 2dlg | Acid Phosphatase A | Hydrolase | *Francisella tularensis* | 15.8 | 3.1 | 221 | 481 | 15 | Unknown |
| 3e2d | alkaline phosphatase | Hydrolase | *Vibrio* | 15.7 | 3.1 | 217 | 502 | 14 | $Zn^{2+}$, $Mg^{2+}$ |
| 2zkt | phosphoglycerate mutase | ISOMERASE | *Pyrococcus horikoshii* | 15.8 | 2.8 | 204 | 381 | 19 | $Zn^{2+}$, $Ca^{2+}$ |

**Supplemental Table 2: DALI comparison of C-terminus of YejM (501-586) to PDB database**

| PDB | Protein name | Z score | rmsd | lali | nres | %id |
|---|---|---|---|---|---|---|
| 6j66 | Chondroitin sulfate/dermatan sulfate endolytic 4-O-sulfatase | 5.2 | 2.9 | 79 | 476 | 9 |
| 1a87 | COLICIN N; | 5.0 | 3.1 | 62 | 297 | 8 |
| 4o56 | SERINE/THREONINE-PROTEIN KINASE PLK1; | 4.9 | 6.3 | 64 | 244 | 11 |
| 5lhx | SERINE/THREONINE-PROTEIN KINASE PLK4; | 4.8 | 2.9 | 60 | 87 | 5 |
| 2n19 | SERINE/THREONINE-PROTEIN KINASE PLK4; | 4.6 | 4.0 | 60 | 87 | 8 |
| 5jov | ALPHA-XYLOSIDASE BOGH31A; | 4.5 | 5.7 | 53 | 949 | 15 |
| 4nkb | PROBABLE SERINE/THREONINE-PROTEIN KINASE ZYG-1; | 3.9 | 3.4 | 60 | 204 | 2 |
| 5wlz | DNA REPAIR PROTEIN XRCC4, MYOSIN-7; | 3.9 | 4.1 | 59 | 206 | 3 |
| 4cys | ARYLSULFATASE; | 3.8 | 3.8 | 77 | 534 | 12 |
| 6hr5 | ALPHA-L-RHAMNOSIDASE/SULFATASE (GH78); | 3.8 | 3.0 | 81 | 376 | 5 |
| 4n7v | SERINE/THREONINE-PROTEIN KINASE PLK4; | 3.7 | 2.5 | 58 | 222 | 5 |
| 4upl | SULFATASE FAMILY PROTEIN; | 3.7 | 3.0 | 83 | 555 | 7 |
| 6dhx | TIPC2; | 3.7 | 2.7 | 68 | 182 | 4 |
| 6qhg | POLYMERASE; | 3.6 | 2.6 | 52 | 119 | 8 |
| 1p49 | STERYL-SULFATASE; | 3.6 | 3.5 | 82 | 549 | 7 |
| 5xq3 | PCRGLX PROTEIN; | 3.4 | 7.1 | 55 | 901 | 16 |
| 4bxr | CPAP; | 3.4 | 3.4 | 53 | 183 | 6 |
| 2obd | CHOLESTERYL ESTER TRANSFER PROTEIN; | 3.3 | 5.9 | 60 | 472 | 10 |
| 5wd6 | SHORT PALATE, LUNG AND NASAL EPITHELIUM CARCINOMA | 3.1 | 3.5 | 65 | 195 | 11 |
| 3cbt | PHOSPHATASE SC4828; | 3.1 | 4.1 | 62 | 210 | 5 |
| 4qtq | XAC2610 PROTEIN; | 3.1 | 3.5 | 57 | 209 | 12 |
| 5hv1 | PHOSPHOENOLPYRUVATE SYNTHASE; | 3.1 | 4.1 | 68 | 847 | 6 |
| 2x2h | ALPHA-1,4-GLUCAN LYASE ISOZYME 1; | 3.0 | 5.0 | 47 | 1025 | 11 |
| 2j6h | GLUCOSAMINE-FRUCTOSE-6-PHOSPHATE AMINOTRANSFERASE | 3.0 | 4.5 | 59 | 609 | 8 |
| 6j9f | DNA-DIRECTED RNA POLYMERASE SUBUNIT ALPHA; | 3.0 | 8.3 | 62 | 1296 | 11 |
| 5mus | L PROTEIN; | 3.0 | 6.9 | 59 | 328 | 12 |
| 3dcz | PUTATIVE RNFG SUBUNIT OF ELECTRON TRANSPORT COMPL | 3.0 | 2.4 | 52 | 170 | 2 |
| 5y83 | MEMBRANE PROTEIN INSERTASE YIDC; | 3.0 | 2.1 | 43 | 342 | 12 |
| 1f34 | PEPSIN A; | 3.0 | 4.9 | 54 | 138 | 2 |

**Supplemental Movies**

Movie 1 file name: SMovie1.mov
Morph between chain C and D of YejM the structure with six monomers in the asymmetric unit. Both chains represent the most “closed” or “open” conformation of the lid.

Movie 2 file name: SMovie2.mov
Same morph as in movie 1, this movie highlights the movement of aromatic residues (red) located at layers II and III, and one aromatic residue (blue) located between layer I and II.
